# A brainstem circuit controls cough-like airway defensive behaviors in mice

**DOI:** 10.1101/2024.09.08.611924

**Authors:** Xiaoshan Xu, Xiupeng Nie, Weijia Zhang, He-Hai Jiang, Bingyi Liu, Yanyan Ren, Tingting Wang, Xiang Xu, Jing Yang, Fujun Luo

## Abstract

The respiratory tract is subject to complex neural control for eupneic breathing and distinct airway defensive reflexes. Growing evidence has highlighted significant heterogeneity of airway-innervating vagal sensory neurons in mediating various respiratory functions, however, the central neuronal pathways and neural circuits involved in the airway regulation remain less understood. Combining whole-body plethysmography (WBP), audio, and video tracking to access breathing and airway defensive behaviors in conscious animals, we developed a quantitative paradigm implementing the mouse as a model to study cough-like defensive behaviors. Using TRAP2 transgenic mice and in vivo fiber photometry, we found that the neural activity in the caudal spinal trigeminal nucleus (SP5C) is strongly correlated with tussigen-evoked cough-like responses. Impairing synaptic outputs or chemogenetic inhibition of the SP5C effectively abolished these cough-like reflexes. Optogenetic stimulation of SP5C excitatory neurons or their projections to the ventral respiratory group (VRG) triggered robust cough-like behaviors without tussive stimuli. Notably, tonic elevation of SP5C excitability caused spontaneous cough-like activities chronically in mice. Together, our data provide strong evidence for a previously unrecognized brainstem circuit that controls cough-like defensive behaviors in mice.

## Introduction

Breathing is essential for maintaining homeostasis of O_2_ and CO_2_ in mammals and is subject to precise neural control. Various innate defensive reflexes, such as apnea, sneezing, coughing, and expiratory reflexes, are crucial for protecting the airways from harmful pathogens and irritants. Dysregulation of the airway-to-brain axis can lead to prevalent and severe clinical problems, including dysphagia, aspiration pneumonia, asthma, chronic cough, and sudden infant death syndrome.

The cough reflex, a crucial airway defensive behavior, is typically elicited by peripheral stimulation of vagal sensory nerve fibers innervating the laryngeal and tracheobronchial airways(Canning, 2006; Canning et al., 2014; Coleridge and Coleridge, 1994). Vagal nerves are heterogeneous in diameter, myelination, conduction velocity, stimuli sensitivity, and physiological functions (Mazzone and Undem, 2016). Strong evidence demonstrates that activation of either mechanical Aδ or chemical-sensitive C-fibers can trigger the cough reflex in various mammalian species(Canning et al., 2004; Widdicombe, 1995). Recent studies have further revealed distinct molecularly-defined subtypes of vagal neurons mediating different airway reflexes, including cough (Chang et al., 2015; Gannot et al., 2024; Kupari et al., 2019; Park et al., 2024; Prescott et al., 2020).

The central fibers of vagal sensory neurons may differentially terminate on second-order neurons in the nucleus of tractus solitarius (NTS) or the paratrigeminal nucleus (Pa5) (Kim et al., 2020; McGovern et al., 2015; Su et al., 2022). These second-order neurons are proposed to transmit cough signals, directly or indirectly, to the ventral brainstem respiratory network, thereby reconfiguring its eupneic function to produce cough motor patterns. The ventral brainstem respiratory network, mainly consisting of the retrotrapezoid nucleus/parafacial respiratory group (RTN/pFRG), the Bötzinger complex (BötC), the pre-Bötzinger complex (preBötC), the rostral ventral respiratory group (rVRG) and the caudal ventral respiratory group (cVRG), has been extensively characterized for its role in coordinating the breathing cycles of inspiration, post-inspiration, and expiration (Del Negro et al., 2018; Krohn et al., 2023; Smith et al., 2013). This remarkably multifunctional network dynamically adapts to control respiration and many non-respiratory motor behaviors such as vocalization, swallowing, coughing, vomiting, and sneezing(Bianchi and Gestreau, 2009; Krohn *et al*., 2023; Li et al., 2021). For example, studies have shown that the cVRG plays crucial roles in many expiration-related behaviors including vocalization(Park *et al*., 2024), swallowing(Subramanian and Holstege, 2009), and coughing(Mutolo, 2017). Indeed, the cVRG has long been proposed as “the cough center” based on various evidence such that pharmacological or electrical stimulation strongly evokes cough in anesthetized cat or rabbit(Bianchi and Gestreau, 2009; Cinelli et al., 2020; Mutolo, 2017). Consistently, pharmacological blockade of glutamatergic transmission in the cVRG completely suppresses cough induced by mechanical stimulation of the tracheobronchial tree in anaesthetized rabbits(Bongianni et al., 2005). Interestingly, however, a recent study using mice as an animal model has elegantly shown that Tac1 neurons in the NTS play a central role in cough-like behaviors by coordinating distinct medullary nuclei including the cVRG to elicit the sequential motor patterns of cough (Gannot *et al*., 2024).

Despite these advances, many key questions remain unanswered: what is the exact function of ventral brainstem respiratory network in the cough reflex? How are distinct nuclei activated by second-order cough relay neurons in the NTS or Pa5, either directly or indirectly? Are there other neuronal populations controlling cough? How are these populations connected with the respiratory network to reconfigure its function? What role might they play in cough hypersensitivity? To address these questions, we utilized the mouse as a powerful model for circuit analysis, employing whole-body plethysmography (WBP), optogenetic, chemogenetic, virus-based tracing and manipulation, and in vivo fiber photometry to explore brain mechanisms of cough and cough hypersensitivity.

## Results

### The SP5C nucleus is necessary for tussigen-evoked cough-like responses in mice

Extensive cough studies have utilized human and larger mammalian models like cats, dogs, rabbits, and guinea pigs, providing critical insights into the mechanisms of cough production and regulation(Bolser, 2004; Canning, 2008; Mutolo, 2017). Although the suitability of mice for cough studies remains debated (Bolser, 2004), their use has recently increased (Chen et al., 2022; Gannot *et al*., 2024; Zhang et al., 2017).

To ensure precise and consistent monitoring of cough-like behaviors in mice, we combined WBP with audio and video tracking of animal behaviors (Figure 1A). We nebulized several widely-used tussigens, including citric acid (CA), capsaicin or irritant gas ammonia (NH3)(Canning *et al*., 2014; Canning *et al*., 2004; Mazzone and Undem, 2016; Zhang *et al*., 2017) to induce the cough reflex. Exposure to these stimuli consistently induced marked cough-like responses in mice, identifiable by tightly-correlated sharp airflow and characteristic cough sound (Figure 1b-e and Figure S1) in wild-type (WT) mice. Cough-like events were rarely observed at baseline or after nebulizing saline (Ctrl), underscoring the specific effectiveness of these tussigens (Figures 1B and 1D). All tussive stimuli reliably triggered cough-like responses in mice, though with different time course. Nebulization of CA or capsaicin rapidly triggered cough-like responses with short latency (< 1 min), which decreased and stopped shortly after nebulization. In contrast, NH3 exposure evoked cough-like responses with longer latency (> 1 min), which reached a high plateau and persisted after nebulization. Notably, NH3-evoked cough-like events were several fold higher in number than those evoked by capsaicin and CA (Figure 1E), likely because NH3 may activate TRPV1, TRPA1 and other target receptors on the sensory neurons(Dhaka et al., 2009). Consequently, given the robustness and specificity of NH3-evoked cough(Sun et al., 2019; Zhang *et al*., 2017), we employed NH3 stimuli for most subsequent experiments. The sneeze reflex, distinguished by unique WBP and audio signals(Chen *et al*., 2022; Gannot *et al*., 2024; Li *et al*., 2021), was occasionally observed following exposure to saline, and capsaicin, but not to NH3, and was excluded from further analysis (Figure S1D). Surgical ablation of bilateral anterior ethmoidal nerves (AENs) had little impact on NH3-evoked cough-like responses, further validating that NH3 exposure predominantly induced cough, rather than sneezing.

**Figure 1.**
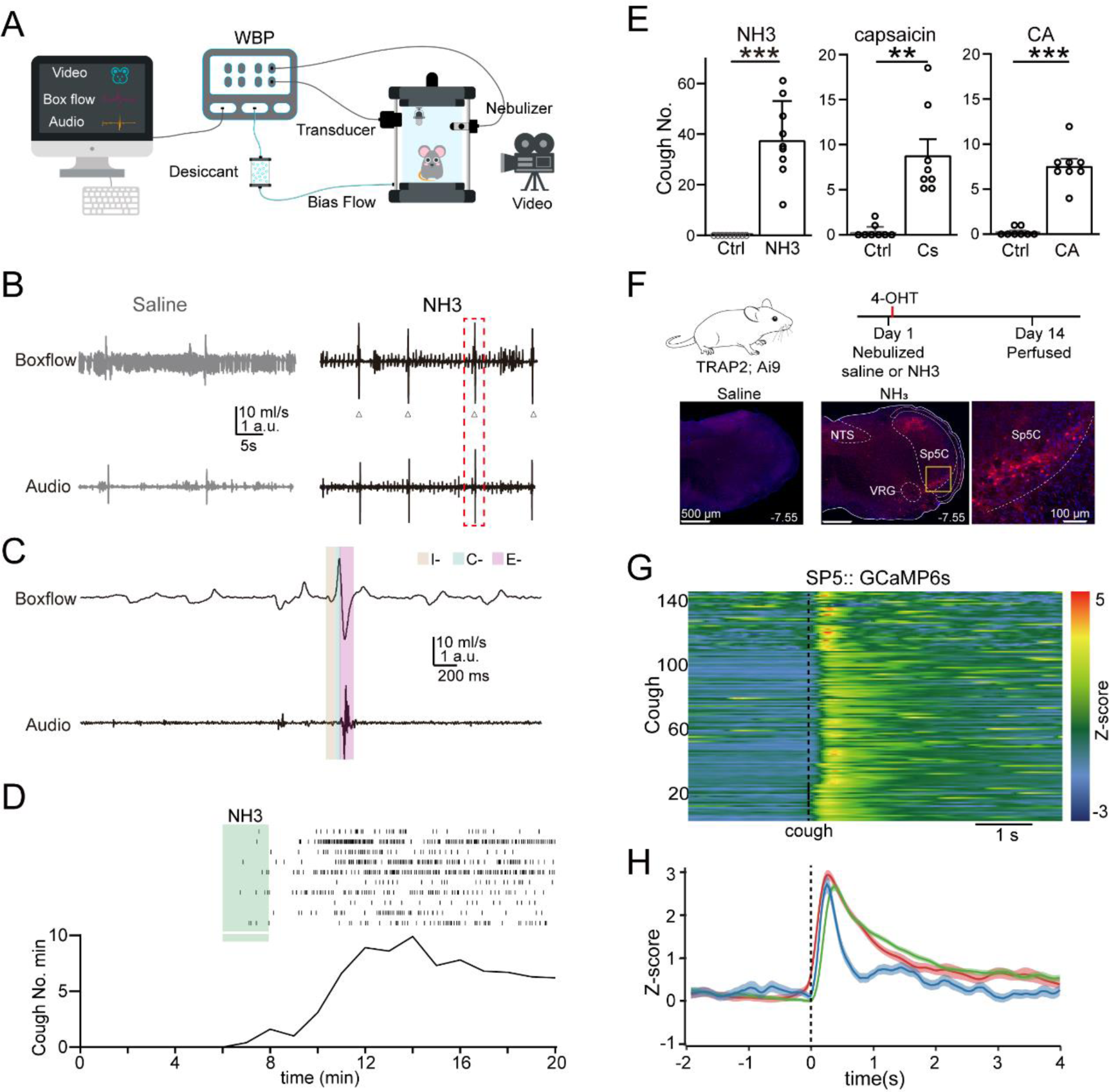
Activation of the SP5C is correlated with tussive-evoked cough responses in mice. (A) Diagram illustrating the setup for cough monitoring in awake mice. (B) Representative traces of box flow and audio signals after exposure to nebulized saline (left) or 1% NH3 (right) in WT mice. Triangles indicate cough responses. (C) Expanded traces of box flow and audio signals from (B) (highlighted by red dashed rectangle), showing eupnoea and a cough along with its corresponding audio signal. I-inspiration; C-compression; E-expiration. (D) Top, raster plot of coughs of individual mice exposed to nebulized NH3 (green bar). Bottom, the number of cough per minute. n = 10 mice. (E) Summary of the number of coughs evoked by various tussigenic stimuli. (F) Experimental strategy and representative images of TRAP2 labeling brainstem neurons during NH3 or saline nebulization. (G) NH3 exposure evoked strong cough-related neuronal Ca^2+^ signals in the SP5C as measured by in vivo fiber photometry. Individual trials of Ca^2+^ signals registered to the peak airflow (time = 0, dashed line) during cough. H the average Ca^2+^ signals in the SP5C of individual mice. n = 3.

To identify brainstem nuclei mediating cough-like behaviors in mice, we genetically labeled activated neurons using TRAP2:: Ai9 mice(Allen et al., 2017; DeNardo et al., 2019) by injecting 4-hydroxytamoxifen (4-OHT) following nebulized saline or NH3 (Figure 1F). Compared to saline, NH3 evoked robust neuronal activation, particularly in the brainstem, including the NTS, SP5, VRG, ventrolateral periaqueductal gray (vlPAG), and the parabrachial nucleus (PBN) (Figure S2 and Figure 1F).

To directly validate whether these identified nuclei are functionally correlated with cough, we performed calcium imaging using in vivo fiber photometry. We injected adeno-associated virus encoding the calcium indicator GCaMP6s (AAV-hSyn-GCamp6s) into various nuclei, including the NTS, VRG, or SP5C, and implanted optic fibers above these regions. Consistent with their established role in cough, real-time neuronal activity in the NTS and VRG were tightly cough-correlated (Figure S2B and S2C). Surprisingly, we also observed a strong correlation between SP5C neuronal activity and cough (Figures 1G and 1H), suggesting that the SP5C nucleus may play an important role in mediating cough.

To test the hypothesis that the SP5C plays a crucial role in mediating cough-like behaviors, we first investigated its necessity for triggering cough-like resonses in mice by abolishing synaptic outputs from the nucleus using AAVs overexpressing the light chain of tetanus toxin (TeTx)(Xu et al., 2012). Unexpectedly, all mice died, likely due to the disruption of the vital physiological functions governed by these nuclei. To overcome this challenge, we utilized neurexin1/2/3 conditional knockout transgenic mice (Nrxn123 cTKO)(Chen et al., 2017; Luo et al., 2020), which enables a strong but not complete blockade of synaptic outputs by Cre-recombinase. In control mice injected with AAV-EGFP in the SP5C, NH3 induced robust cough-like responses similar to wild-type animals. However, Nrxn123 cTKO mice injected with AAV-Cre-EGFP (TKO) exhibited significantly reduced cough-like responses to NH3 (Figures 2A-C). Using a similar strategy, we found that pan-neurexin ablation in the VRG also blocked NH3-evoked cough-like responses (Figure S3). Therefore, impairing synaptic outputs from either the SP5C or VRG markedly blocked NH3-evoked cough-like responses, suggesting their crucial roles in triggering the cough reflex.

**Figure 2.**
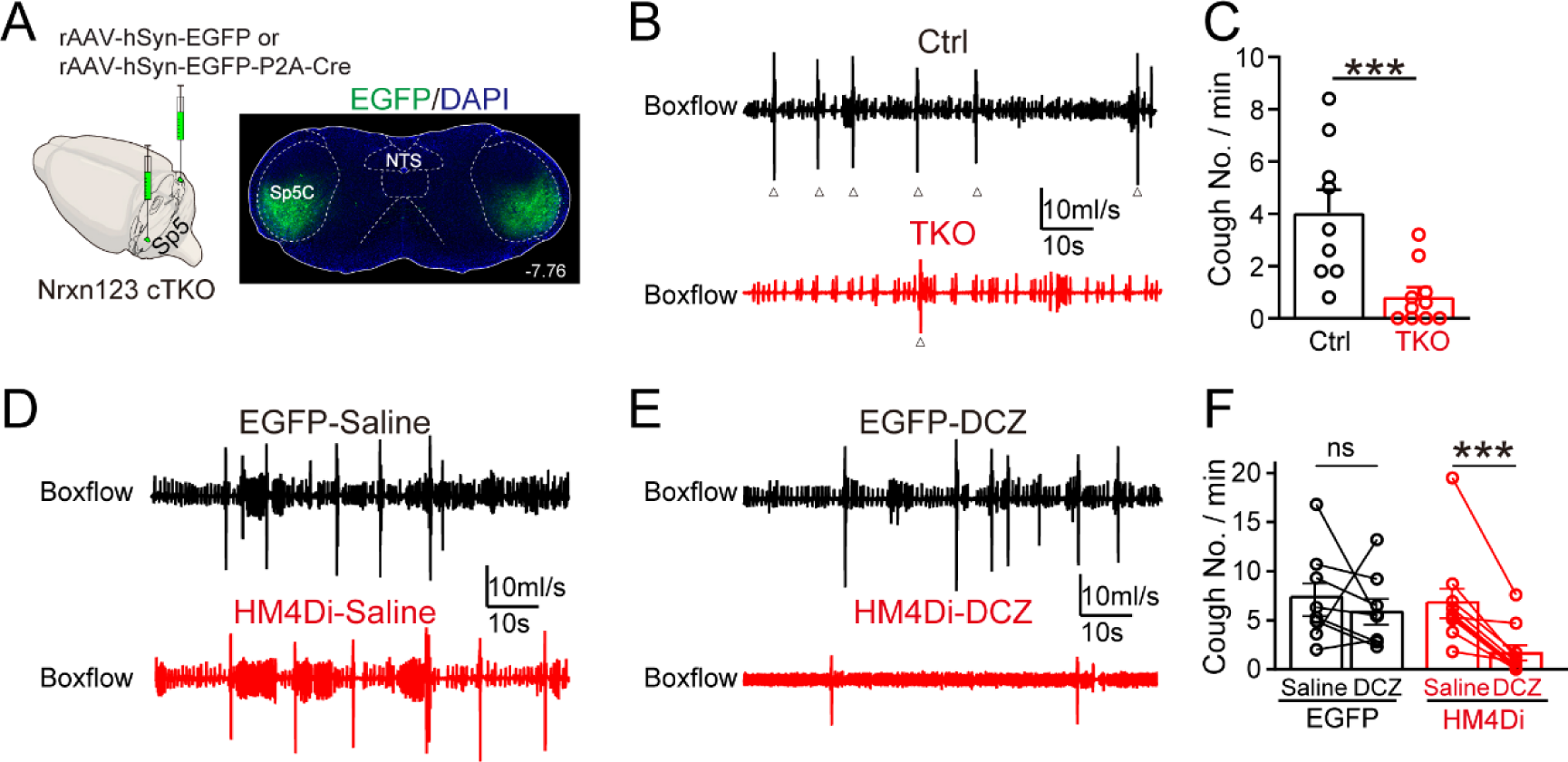
The SP5C is necessary for tussive-evoked cough responses in mice. (A) Experimental strategy, combining stereotactic injection of AAV overexpressing Cre with transgenic mice of neurexin1/2/3 conditional knockout mice, to impair synaptic outputs of the SP5C. (B) Representative traces of box flow signals following exposure to 1% NH3 in mice with or without pan-neurexin deletion in the SP5C. (C) Summary of NH3-evoked cough responses in Ctrl and Nrxn123 TKO in the SP5C. (D-E) Representative traces of box flow signals following exposure to 1% NH3 in mice with EGFP or hM4Di expression in the SP5C when saline (D) or DCZ (E) was injected intraperitoneally. (F) Summary of NH3-evoked cough with and without chemogenetic inhibition of the SP5C. Data are means ± SEM. Statistical difference was assessed by Student’s t-test. (***P < 0.001).

To achieve more timely control of neuronal activity, we used chemogenetic approach by bilaterally injecting AAV-hM4Di-mCherry into the SP5C. Inactivation of SP5C neurons using deschloroclozapine (DCZ)(Nagai et al., 2020) strongly inhibited NH3-evoked cough-like responses in mice (Figures 2D-2F). In contrast, three control groups, either with AAV-mCherry injection or saline treatment, showed robust cough-like activities upon NH3 nebulization (Figures 2D-2F). These data collectively demonstrate the essential function of the SP5C in tussigen-evoked cough-like behaviors in mice.

### Activating SP5C CaMKII^+^ excitatory neurons is sufficient to trigger cough in mice

We then investigated whether activating the SP5C could directly trigger cough-like behaviors using an optogenetic technique. We injected AAV-ChrimsonR, a red-light-sensitive variant of channelrhodopsin(Klapoetke et al., 2014), into the SP5C and implanted an optic fiber (Figure 3A). Optogenetic stimulation of the SP5C caused pronounced motor behaviors that confounded cough response measurement, probably due to concurrent activation of distinct motor control circuits. To address this challenge, we used transgenic Cre-lines for more precise manipulation of specific cell types. We tested the functional impacts of activating excitatory or inhibitory neurons by selectively expressing Cre-dependent ChrimsonR in the SP5C of VGluT2-IRES-Cre or GAD2-IRES-Cre mice. Optogenetic stimulation of VGluT2^+^ or GAD2^+^ neurons caused a remarkable increase or decrease in the respiration rate of mice, respectively (Figures 3B and 3C). Surprisingly, however, activation of neither VGluT2^+^ excitatory neurons nor GAD2^+^ inhibitory neurons triggered cough (Figures 3B and 3C).

**Figure 3.**
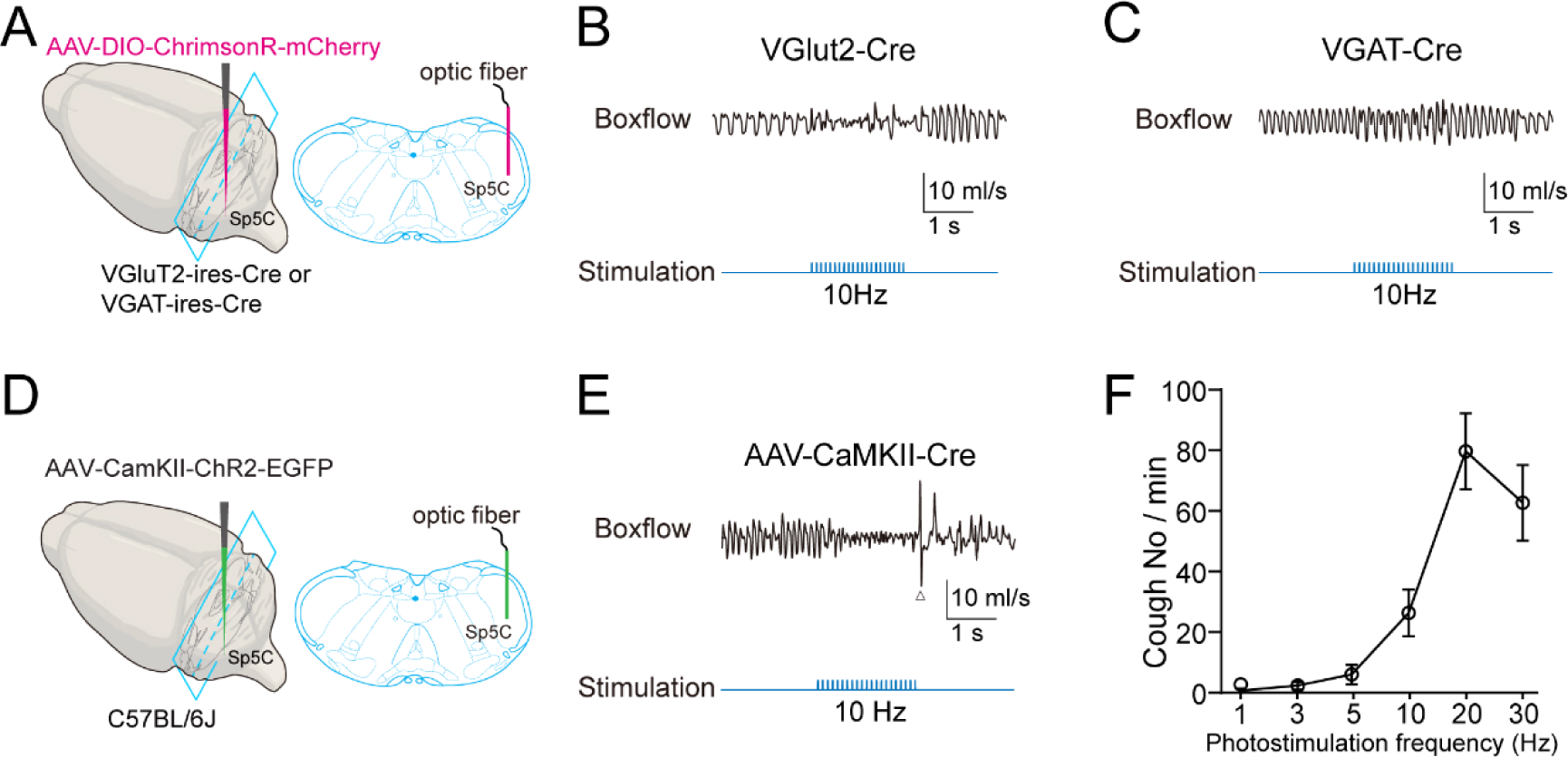
Optogenetic stimulation of SP5C excitatory neurons is sufficient to trigger robust cough activities in mice. (A) Strategy for optogenetic stimulation of specific cell types in the SP5C. (B-C) Representative box flow signals following optogenetic stimulation of VGlut2^+^ excitatory neurons, or VGAT^+^ inhibitory neurons, in the SP5C. (D) Experimental strategy of optogenetic stimulation of CaMKII^+^ excitatory neurons in the SP5C. (E) Representative box flow signals of mice during optogenetic stimulation of CaMKII^+^ neurons in the SP5C. (F) Summary of the number of coughs (converted into the rate in minute) evoked by optogenetic stimulation at various frequencies. Data are means ± SEM. Statistical difference was assessed by Student’s t-test. (**P < 0.01; ***P < 0.001).

Interestingly, when we injected AAV-CaMKII-ChrimsonR into the SP5C, we found that optogenetic stimulation of CaMKII^+^ neurons induced strong cough-like activity in the absence of tussive stimuli (Figures 3D-F). These data suggest that a specific population of excitatory neurons in the SP5C, although their precise identity remains to be further characterized, plays a crucial and sufficient role in triggering cough-like behaviors in mice.

### Activating the SP5C projections to the VRG triggers cough in mice

Next, we investigated how the SP5C neurons connect with the respiratory network to produce cough motor behavior. Given that the VRG is well-known as the cough center (Bolser and Davenport, 2002) and its critical roles in NH3-evoked cough-like responses we observed (Figures S2 and S3), we hypothesized that the SP5C may project to the VRG to induce cough. To test this hypothesis, we characterized the synaptic connectivity between the SP5C and the VRG using anterograde and retrograde transynaptic tracing techniques.

We injected AAV-hSyn-EGFP into the SP5C of WT mice and observed widespread projection of EGFP^+^ afferent fibers to different parts of ventral brainstem, including preBotzinger complex, rVRG, and cVRG (Figures 4 and S4). To complementarily verify that SP5C and VRG neurons are monosynaptically connected, we conducted retrograde transsynaptic tracing using pseudotyped EnvA-G-deleted rabies virus (RV)(Wickersham et al., 2007). Unilateral RV injections into the VRG resulted in reliable labeling of SP5C neurons (Figures 4A and 4B), suggesting monosynaptic projections originating from the SP5C.

**Figure 4.**
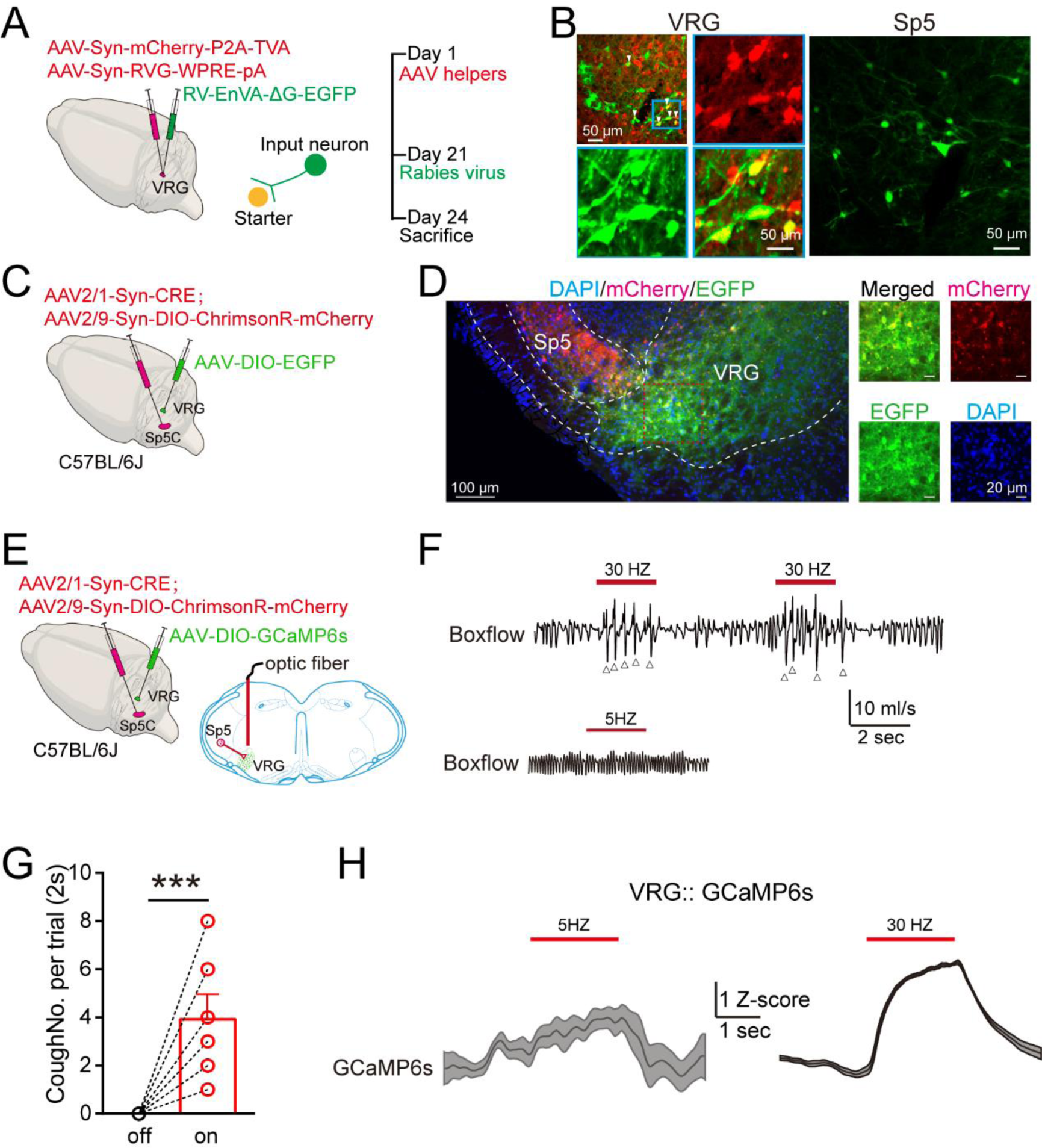
The SP5C projects to the VRG to trigger cough in mice. (A) Schematic and time course of the retrograde tracing strategy. (B) RV labeling of VRG neurons receiving monosynaptic inputs directly from the SP5C. (C) Schematic of the anterograde tracing strategy. (D) AAV2/1 labeling of the SP5C afferent terminals innervating VRG neurons. (E) Strategy of optogenetic stimulation of SP5C projections to the VRG and simultaneous recording of Ca^2+^ signals from VRG neurons. (F) Representative box flow signals during optogenetic stimulation, at either high or low frequency, of the SP5C projections to the VRG. (G) Summary of the number of coughs evoked by optogenetic stimulation. (H) Summary of the GCaMP6s signals evoked by optogenetic stimulation of the SP5C projections to the VRG. Data are means ± SEM. Statistical difference was assessed by Student’s t-test. (***P < 0.001).

To further confirm their direct synaptic connectivity, we used a dual virus strategy. We co-injected AAV2/1-Cre, an anterograde transsynaptic variant, and AAV2/9-DIO-Chrimson-mCherry into the SP5C to label presynaptic neurons. AAV2/9-DIO-EGFP was injected into the VRG to label postsynaptic neurons only if they were synaptically connected with AAV2/1-Cre infected SP5C neurons (Figure 4C). We observed prominent afferent fiber labeled by mCherry coursing from the SP5C to the VRG (Figure 4D). Additionally, numerous EGFP^+^ neurons were reliably observed in the VRG. Moreover, frequent co-labeling of EGFP and mCherry was observed near the soma or proximal dendrites of VRG neurons (Figure 4D). Together, these data provide strong evidence for a monosynaptic connection from the SP5C to the VRG.

To test the functional impact of the SP5C-VRG pathway on the cough-like behaviors in mice, we overexpressed ChrimsonR in the SP5C and implanted the optic fiber above the VRG, where it receives synaptic input from the SP5C (Figure 4E). Remarkably, selective photostimulation of SP5C projections to the VRG triggered robust cough-like behaviors in mice (Figure 4F). Interestingly, low-frequency optogenetic stimulation barely evoked cough-like activities, suggesting that cough can only be triggered during intense stimulation of afferent fibers (Figures 4F and 4G). In acute brainstem slice, we found that SP5C excitatory neurons showed relatively high membrane resistance and fired high-frequency action potentials after current injection above their threshold (Figure S5). In addition, optogenetic stimulations reliably induced action potential firing of SP5C neurons at a wide range of frequencies from 1 Hz to as high as 50 Hz (Figure S5).

In some animals, we injected AAV2/9-DIO-GCaMP6s into the VRG in conjunction with injection of AAV2/1-Cre and AAV2/9-DIO-Chrimson-mCherry into the SP5C while implanting an optic fiber above the VRG. Optogenetic stimulation of the SP5C-VRG projection induced remarkable activity-dependent Ca^2+^ signals in the VRG (Figure 4H). Together, these results demonstrate that specific activation of the SP5C-VRG circuitry is sufficient to trigger robust cough-like activities in mice without tussive stimuli.

### Tonic excitation of the SP5C chronically induces spontaneous cough

Thus far, our findings underscore the critical role of the SP5C in cough production, primarily through CaMKII^+^ excitatory neurons projecting to the VRG. We next investigated whether neural activity in the SP5C contributes to cough hypersensitivity, a hallmark feature manifesting in chronic cough(Chung et al., 2022). To explore this, we elevated SP5C neuronal excitability by overexpressing NaChBac (Figure 5A), a bacterial voltage-gated Na^+^ channel(Ren et al., 2001; Xue et al., 2014).

**Figure 5.**
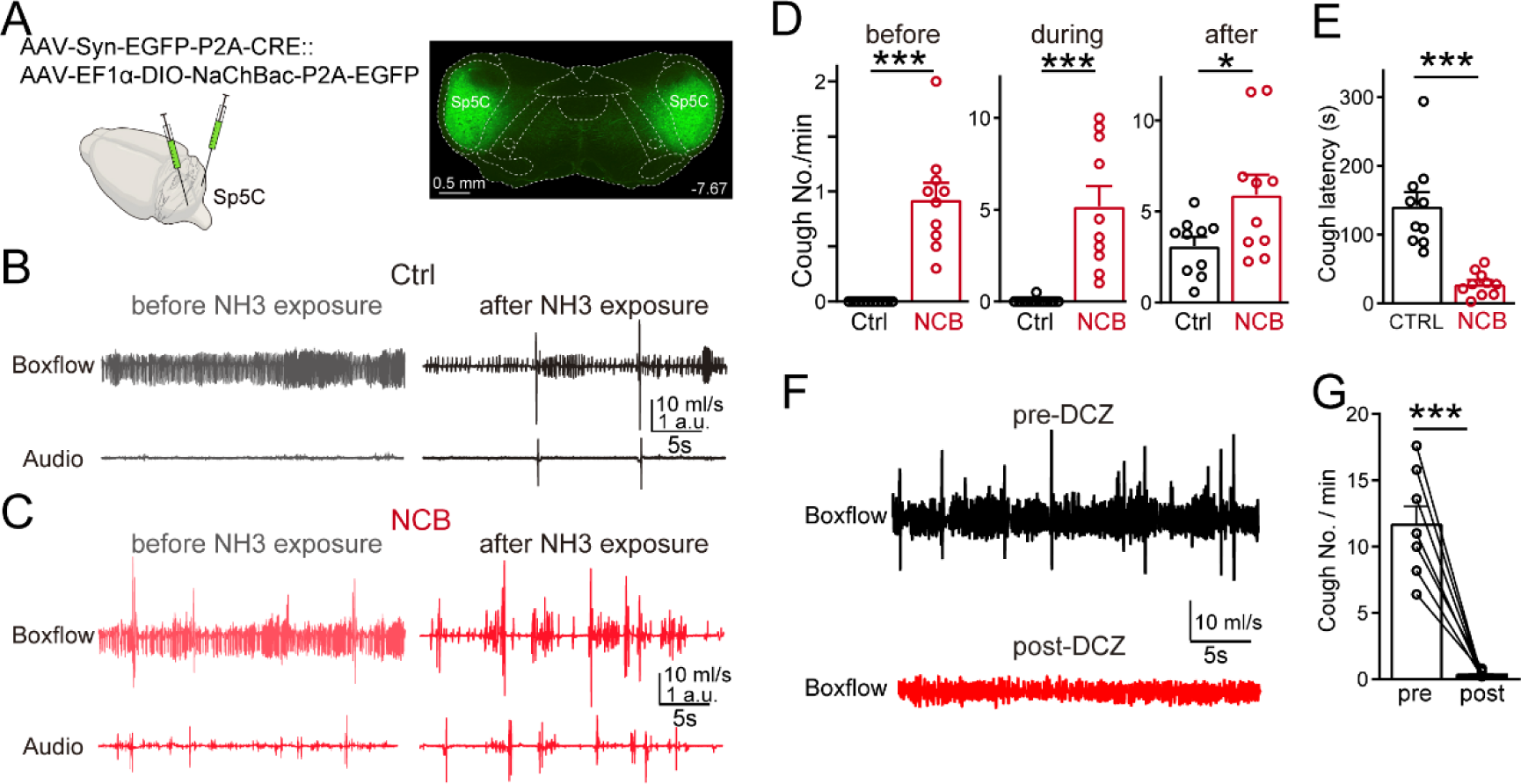
Elevating SP5C neuronal excitability induces spontaneous cough and cough hypersensitivity in mice. (A) Experimental strategy (left) and fluorescence image illustrating overexpression of NaChBac in the SP5C. (B-C) Representative traces of box flow and audio signals before and after exposure to NH3 in mice overexpressing EGFP as control (B) or NaChBac (C) in the SP5C. (D) Summary of the number of coughs per minute before, during and after NH3 exposure. (E) Summary of the cough latency (the time of first cough recorded after NH3 nebulization). (F) Representative traces of box flow signals before and after injection of DCZ in mice overexpressing NaChBac in the SP5C and HM4Di in the cVRG. (G) Summary of the number of spontaneous cough events before and after chemogenetic inhibition of the cVRG. Data are means ± SEM. Statistical difference was assessed by Student’s t-test. (*P < 0.05; ***P < 0.001).

In control mice injected with AAV-EGFP in the SP5C, spontaneous cough-like activity was barely observed (Figure 5B). In contrast, mice injected with AAV-NaChBac exhibited spontaneous cough-like behavior (Figure 5C). We chronically monitored this spontaneous behavior after virus injection. Mice began to display spontaneous cough-like behaviors approximately 3-4 days after AAV inoculation. The frequency increased with prolonged expression of NaChBac, peaked roughly 2-3 days later, and then declined, likely due to decreased mobility and health of mice. When mice began to display spontaneous cough-like behaviors, we exposed them to NH3. Both control and NaChBac-overexpressing mice showed robust cough-like responses to NH3 (Figures 5B-5D). However, the number of NH3-evoked cough-like responses was significantly higher in mice overexpressing NaChBac than in control mice (Figures 5B-5D). Additionally, the ability to suppress cough-like responses in mice with NaChBac overexpression appeared to decrease, as evidenced by the remarkably decreased latency of the first occurrence of cough-like responses during nebulization, while the number of cough-like responses increased compared to control (Figure 5E). To verify that overexpression of NaChBac indeed increases the excitability of SP5C neurons, we recorded their spiking activity in acute brainstem slice using whole-cell patch clamp recording. Indeed, compared to control, NaChBac-overexpressing neurons fired spontaneously (Figure S6). Further, injected currents caused action potential firings at higher frequency and sustained depolarization long after the end of current injection. Interestingly, when the same virus was injected into the Sp5C of TRAP2 mice, NH3 induced more abundant TRAP2-labeled neurons, most of which were overlapped with NaChBac-EGFP, suggesting an enhanced activation of SP5C neurons (Figure S7).

Finally, we examined whether the spontaneous cough-like behaviors observed in NaChBac-overexpressing mice requires the intact function of the VRG. We injected bilaterally AAV-hM4Di-mCherry into the VRG followed by injecting AAV-NaChBac into the SP5C. Inactivation of VRG neurons by injecting DCZ nearly abolished all spontaneous cough-like behaviors (Figure 5F and 5G). Together, these findings demonstrate that elevating neuronal excitability in the SP5C remarkably enhances the cough sensitivity of mice, which can last as long as 48-72 hours. Moreover, albeit through a non-physiological manipulation, we show for the first time that spontaneous cough-like behaviors can occur in an animal model, warranting future study of its mechanism and pharmacological regulation.

## Discussion

Consistent with recent studies(Chen *et al*., 2022; Gannot *et al*., 2024), our present findings provide multiple lines of evidence that mice are a suitable model for studying the central mechanisms of cough behavior, even in the conscious state. Several key observations support this assertion. First, various tussive agents reliably elicit characteristic airflow changes and cough sounds, which can be consistently monitored (Figure 1). Second, tussive stimuli evoke robust neural activity in multiple brain regions tightly correlated with cough in awake animals. Because the physiological states of the animal, i.e. anesthetized *vs.* awake, significantly impact the regulation of the cough reflex, measuring the cough responses in awake animals is crucial, especially for evaluating potential antitussive agents. Third, impairing synaptic outputs or neuronal activity from these nuclei effectively blocks tussigen-evoked cough (Figure 1). Finally, selective activation of these nuclei or pathways can reliably induce cough in the absence of tussigen (Figures 2-4). Consequently, transgenic mouse models hold great promise for advancing the study of genetic and neural mechanisms of cough and cough hypersensitivity.

Utilizing transgenic mice and various circuit-based tools, we have identified a novel brainstem circuit necessary and sufficient for triggering cough in mice. Remarkably, our findings demonstrate that bi-directional regulations of SP5C neuronal activity exert prominent but opposite impacts on cough activity, underscoring its central role in cough production in mice (Figures 1, 2, 3). Accumulating studies have shown that the SP5C may exert diverse physiological roles in the sneezing reflex(Li *et al*., 2021), self-grooming(Xie et al., 2022) and craniofacial nociception(Xue et al., 2024), probably via different neuronal subtypes located in distinct subregions or circuitry of the SP5C. For example, NMDR^+^ neurons are specifically involved in sneezing reflex(Li *et al*., 2021) while Clbn2^+^ neurons are involved in orofacial self-grooming(Xie *et al*., 2022). We have shown that CaMKII^+^ excitatory neurons play an essential role in mediating cough responses (Figure 3), but their precise molecular identity remains to be further characterized.

Our preliminary data suggest that the SP5C does not receive direct synaptic projection from the NTS, but rather from the Pa5, the nucleus known to be specifically activated by jugular sensory neurons. However, we do not yet have conclusive evidence regarding which vagal sensory neurons project tussive information to the Pa5. Strong evidence has recently shown that Tac1^+^ neurons in the NTS directly project to the cVRG and contribute to the cough-like behavior in mice (Gannot *et al*., 2024). It is likely that multiple neuronal pathways are involved in eliciting the cough reflex. Future studies are needed to explore whether these pathways function independently or interactively in cough control. Our data also suggest that the SP5C does not receive direct synaptic projection from the NTS, but rather from the Pa5 and other nuclei and cortical areas, such as the red nucleus, S1, and M1 cortex. It is crucial to investigate the specific contributions of distinct circuits to cough, particularly cough hypersensitivity.

Recent studies have revealed a high prevalence of chronic cough associated with RFC1 polymorphism of nucleotide repeat expansion in CANVAS patients(Cortese et al., 2019). This polymorphism has been identified in up to 25% of patients with unexplained or refractory chronic cough (UCC/RCC)(Guilleminault et al., 2023; Turner et al., 2023), indicating a strong genetic link with UCC/RCC. Therefore, it is of great interest to exploit the myriad advantages of transgenic mice to study the genetic and neural mechanisms of UCC/RCC in the future.

## Acknowledgments

We thank Peng Cao, Bo Zhang, and Thomas Sudhof for their insightful advice on the project. The study was supported by Major Project of Guangzhou National Laboratory #GZNL2024A02001 (F.L.).

## Author information

These authors contributed equally: Xiaoshan Xu, Xiupeng Nie

## Author contributions

X. X., X.N., F.L. conceived the project. X. X., X.N., T.W., W.Z., H.J., B.L., Y.R. collected the data and performed the experiments. X. X., X.N., T.W., W.Z., H.J., F.L. analyzed the data. F.L. wrote the original draft. X.X. and F.L. revised, edited and approved the final version. F. L. supervised the project.

## Methods

### Animals

All experiments were conducted in accordance with the guidelines established by the Institutional Animal Care and Use Committee at Guangzhou National Laboratory. Male mice were utilized for most experiments except specified otherwise. The mice were housed under standard laboratory conditions, with a temperature range of room temperature and 40-60% humidity maintained, and subjected to a 12-hour light-dark cycle (lights on from 7:00 to 19:00) with food and water freely available. No statistical tests were used to predetermine the sample size. All experiments were conducted blindly, with the experimenter unaware of the mouse genotypes or adeno-associated viral vectors (AAVs) used. The experiments were conducted using mice aged between two to three months. C57BL/6J wild-type mice were used. All transgenic mouse lines, including Nrxn123 cKO(Chen *et al*., 2017), VGluT2-IRES-Cre (JAX#: 028863)(Vong et al., 2011), VGAT-IRES-Cre (JAX#: 028862)(Vong *et al*., 2011), and FosTRAP2(JAX#: 030323)(Allen *et al*., 2017) were previously reported.

### Virus vectors

All AAV vectors were purchased from Brain VTA (Wuhan, China). The final titers of viral vectors were diluted specifically in the range from 3−8×10^12^ particles/ml before use. Rabies viruses were purchased from Brain VTA (Wuhan, China). Pseudo rabies viruses were purchased from Brain Case (Wuhan, China).

### Stereotactic injections of AAVs

Mice were anesthetized via intraperitoneal injection of chloral hydrate (0.5 mg/kg) (Sangon Biotech, Shanghai, China). Following anesthesia, standard surgical techniques were employed to expose the brain areas above the brainstem. Coordinates for virus injections were determined relative to the Bregma. For the NTS, the coordinates were set at Anteroposterior (AP): −7.48 mm, Mediolateral (ML): ± 0.45 mm, and Dorsoventral (DV): −4.4 mm. For the SP5C, the coordinates were AP: −7.83 mm, ML: ± 1.85 mm, DV: −5.4 mm. For the VRG, the coordinates were AP: −7.48 mm, ML: ± 1.25 mm, DV: −5.5 mm. The AAVs were injected into the designated brainstem nuclei using a glass pipette connected to a microinjector system Nanoject III (Drummond Scientific Company, Pennsylvania, USA). The injection was performed at a controlled flow rate of 60 nl/min. The volume of AAV injected was determined based on experimental requirements and viral titration. The injected volume was carefully calibrated to ensure precise delivery into the target nuclei. Experiments were conducted at least 2 weeks after virus injections to allow for sufficient viral expression and integration into the target cells.

### Optogenetic and in vivo fiber photometry

After viral injections, a ceramic ferrule equipped with an optic fiber (200 μm diameter and numerical aperture (N.A.) of 0.37 for optogenetics and fiber photometry) was implanted at the target brain regions, namely the NTS, SP5C, and VRG. The fiber tip was meticulously positioned over the target nuclei, ensuring precise placement. Secure fixation of the ferrule to the skull was achieved using dental cement, ensuring stability throughout the experiment. Following implantation, the surgical site was sutured, and topical antibiotics were administered to minimize the risk of inflammation and infection.

Optogenetic and fiber photometry experiments were conducted at least 3 weeks after fiber implantation to allow for adequate viral expression and integration. After experiment, each animal underwent histological analysis to verify virus expression and fiber localization. Only mice demonstrating appropriate viral infection and accurate fiber implantation at the target nuclei were included in subsequent data analysis.

### Chemogenetic inhibition

To chemogenetically silence the SP5C nucleus, AAV-hM4Di-mCherry was bilaterally injected into the SP5C of WT mice. After 3 weeks of viral injection, DCZ solution (0.1 mg/ml in saline, 0.1mg/Kg) was intraperitoneally injected into mice. Saline was used as a control. Cough tests were performed after 30 mins.

### Breathing and cough measurement

After a recovery period of 3 weeks following surgery, mice were acclimated for three consecutive days being handled daily by experimenters to minimize stress. On the day of the experiment, mice were transferred to the testing room and habituated to the experimental chamber for 1 hour before the commencement of tests.

Whole-body plethysmography (TOW Tech, Shanghai, China) was utilized to monitor animal breathing and cough responses. This technique enabled continuing measurement of respiratory parameters, including the timing and magnitude of inspiratory and expiratory efforts. Throughout the experiments, mice were awake and freely moving within the recording chamber.

High-resolution audio and video recordings of the animals’ movements were captured using a microphone (Boya, Shenzhen, China) and video camera (Logitech, Shanghai, China). Pressure changes induced by respiration within the recording chamber were continuously monitored before and after the administration of tussive stimuli.

A nebulized solution (1 mL) containing capsaicin (10 μM), CA (0.1 M), NH3 (0.1%), or saline was introduced into the recording chamber at a constant speed of 0.5 ml/min. All signals were simultaneously recorded for 5 min before, 2 min during, and 10 min after each nebulization event. A minimum interval of 30 min was allowed for full recovery between consecutive challenges.

Cough responses were characterized by rapid timing (<0.1 s for the entire maneuver) and a significant magnitude of inspiratory and expiratory efforts (>500% of expiratory pressure during tidal breathing). This comprehensive approach allowed for the accurate assessment and analysis of cough responses in awake mice.

### Fiber photometry recording

An AAV expressing the genetically encoded calcium indicator GCaMP6s was injected into the targeted nucleus using stereotactic techniques. Subsequently, a fiber optic was implanted 100 μm above the injected region to facilitate optical recordings. A fiber photometry recording system (Inper, Hangzhou, China) was utilized to both activate and record calcium-related fluorescence transients from different nuclei in freely moving mice. Prior to each experiment, the light intensity of the optic fiber was carefully adjusted to ensure optimal fluorescence signal detection, typically within a range of 40 μW. Following exposure to tussive stimuli or optogenetic activation of presynaptic neurons, cough-associated fluorescence signals were acquired in real time from awake mice. Data acquisition was synchronized with coughing episodes to precisely capture neural activity changes associated with cough events. Post hoc analysis of the recorded fluorescence data was conducted to extract calcium transients corresponding to neural activity changes in the target brain regions. Signals were normalized and expressed in Z-score to facilitate comparison across experimental conditions and between animals.

### Acute brain slice for electrophysiology

Coronal brainstem slices were prepared similarly as described previously (Luo *et al*., 2020). In brief, mice of 8-12 weeks were decapitated; brains were rapidly isolated and fixed on the cutting platform a vibratome (VT1200s; Leica, USA), which was immersed in oxygenated cold ACSF containing (in mM): 119 NaCl, 26 NaHCO3, 10 glucose, 1.25 NaH2PO4, 2.5 KCl, 0.05 CaCl2, 3 MgCl2, 2 Na-pyruvate, and 0.5 ascorbic acid, pH 7.4. Transverse slices of 250 μm were sectioned and transferred into a beaker with bubbled ACSF containing (in mM): 119 NaCl, 26 NaHCO3, 10 glucose, 1.25 NaH2PO4, 2.5 KCl, 2 CaCl2, 1 MgCl2, 2 Na-pyruvate, and 0.5 ascorbic acid, pH 7.4. Slices were recovered at 35°C for 30, and stored at room temperature (∼21-23°C) for experiments.

### Whole-cell patch-clamp recordings

Patch-clamp recordings were made from visually identified SP5C neurons using an Axopatch 700B amplifier (Molecular Devices, USA) and pClamp 11 software (Molecular Devices, USA). Patch pipettes were pulled from borosilicate glass capillaries using a two-stage vertical puller (PC-100, Narashige, Japan). Pipette resistance was 4-6 MΩ when filled with an internal solution containing (in mM): 145 K-gluconate, 6 KCl, 10 HEPES, 3 Na2-phosphocreatine, 4 Mg-ATP, 0.3 Na2-GTP, 0.2 EGTA, 2 Qx-314; pH 7.2 with KOH. For optogenetics, a light simulation of 5 ms was delivered by LED (pE-300 white, CoolLED, UK). Action potentials of SP5C neurons were recorded by holding the membrane potential at approximately −70 mV. For optogenetics, a light simulation of 5 ms was delivered by LED (pE-300 white, CoolLED, UK).

### Histology and immunohistochemistry

Mice were anesthetized and perfused at speed of 1ml/min with 1×PBS for 10 min followed by 4% paraformaldehyde (PFA) for 10 min. Brains were removed and stored in 4% PFA overnight at 4C. Brain tissues were cryo-protected with 50% sucrose for 48 hours and then embedded in OCT (Cat#: 14020108926, Leica) and rapidly frozen on dry ice. Transverse sections at 30 μm were cut at −20 °C using a cryostat (CM3050-S, Leica) and mounted directly on Superfrost Plus slides (Cat#: PRO-04, Matsunami Platinum Pro). Samples were mounted with a Vectashield hard-set antifade mounting medium (Cat#: H-1500, Vector Laboratories). Images were acquired using Olympus slide scanner or Zeiss LSM800 confocal microscopy with a 63x oil-immersion objective (1.4 numerical aperture) and analyzed in Zeiss 2.6 and Image J software.

### Statistical analysis

Statistical analyses are described in each corresponding experimental figures. The number of samples is indicated in figures and figure legends.

## Data availability

All data supporting the findings of this study are available from the corresponding author (F.L.).

---

**Figure 1-Figure supplement 1.**
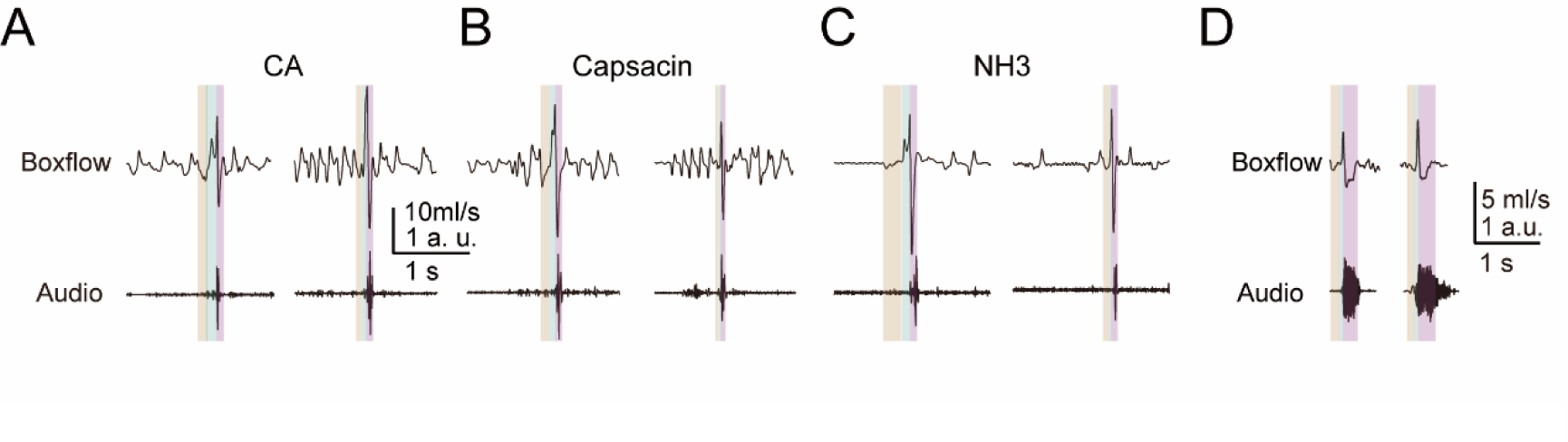
Cough responses with variation in the boxflow waveforms can be observed across different tussive stimuli. (A) Representative box flow and audio signals of two cough events evoked by nebulized CA (0.1M). (B) Representative box flow and audio signals of two cough events evoked by nebulized capsacin (10 µM). (C) Representative box flow and audio signals of two cough events evoked by nebulized NH3 (1%). (D) Representative box flow and audio signals of two sneeze events evoked by nebulized capsacin (10 µM).

**Figure 1-Figure supplement 2.**
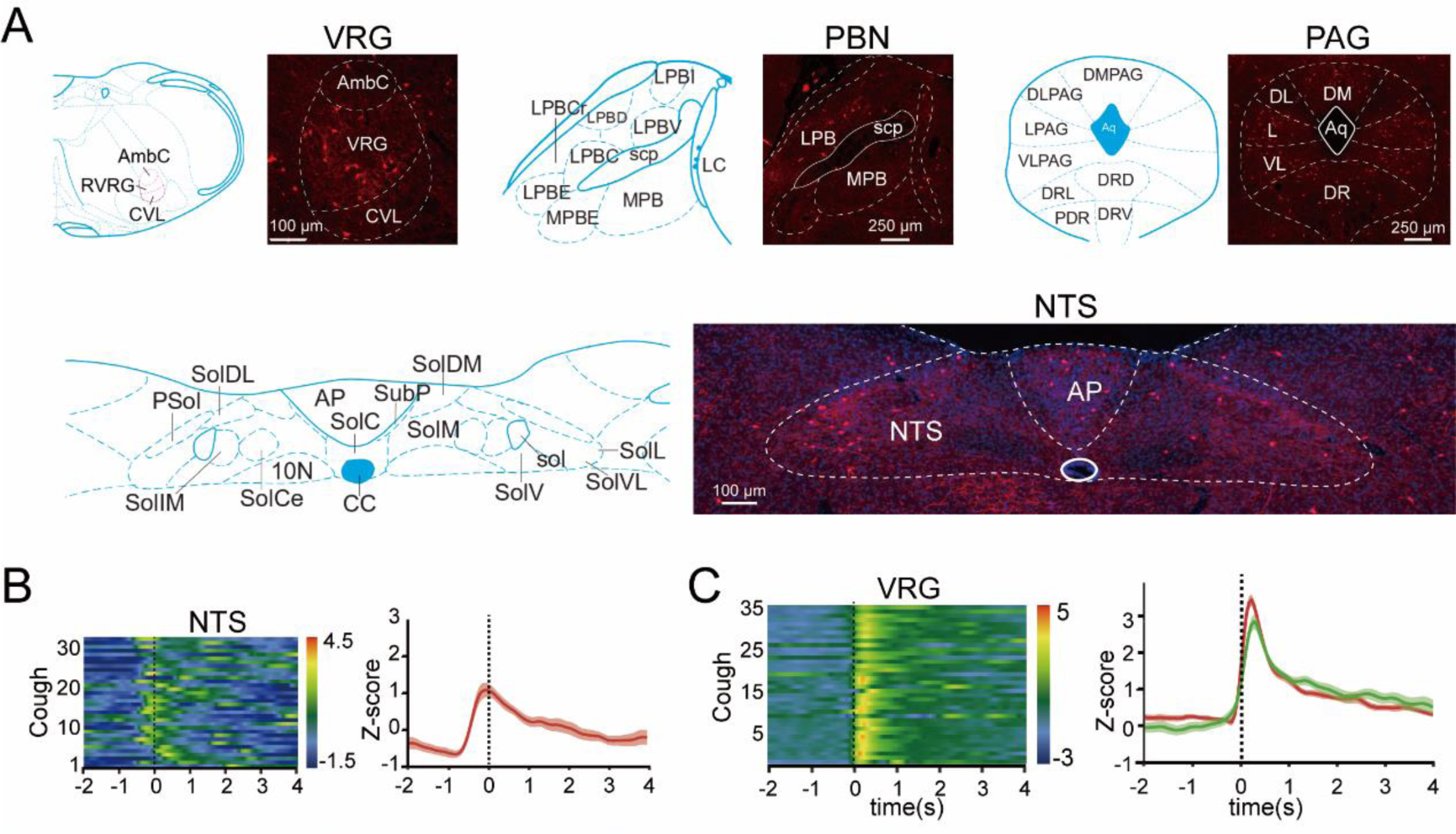
The neural activity in the NTS or VRG is correlated with tussive-evoked cough responses in mices. (A) Representative images of TRAP2 labelled neurons in the VRG, PBN, PAG, and the NTS following NH3 exposure in TRAP2::Ai9 mice. (B) Individual (left) and average (right) cough-correltaed GCaMP6s signals in the NTS recorded by in vivo fiber photometry. (C) Individual (left) and average (right) cough-correltaed GCaMP6s signals in the VRG recorded by in vivo fiber photometry.

**Figure 2-Figure supplement 1.**
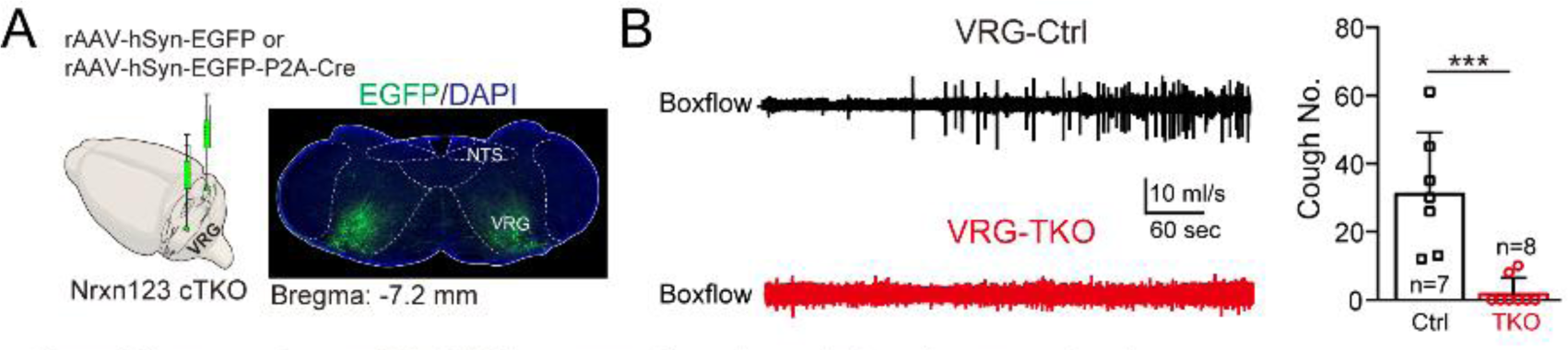
The VRG is necessary for tussive-evoked cough responses in mices. (A)Strategy and fluorescence image showing AAV mediated pan-neurexin ablation in the VRG. (C) Representative box flow signals of mice without or with pan-neurexin deletion in the VRG following NH3 exposure (left) and the summary of cough number (right).

**Figure 4-Figure supplement 1.**
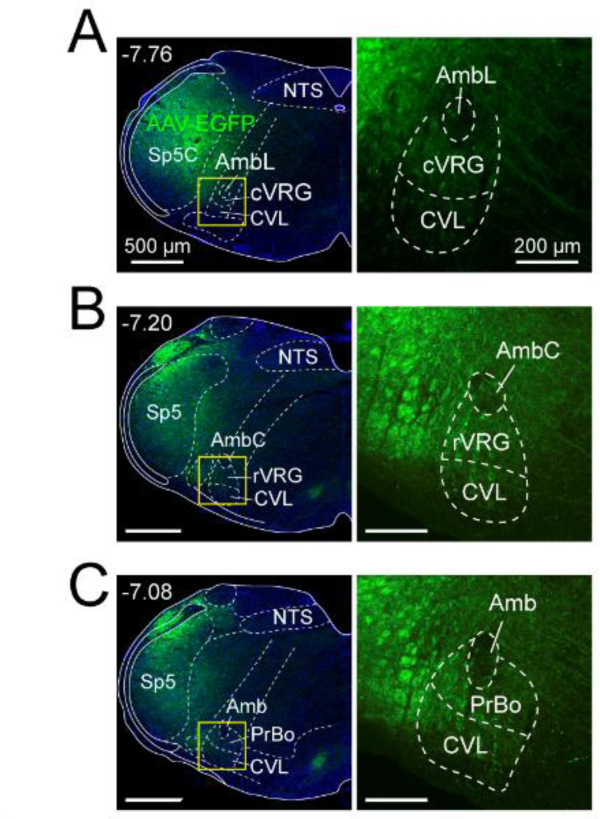
The SP5C neurons project their terminals to the respiratory network, including the cVRG (A), rVRG (B) and preBotzinger complex (PrBo) (C).

**Figure 5-Figure supplement 1.**
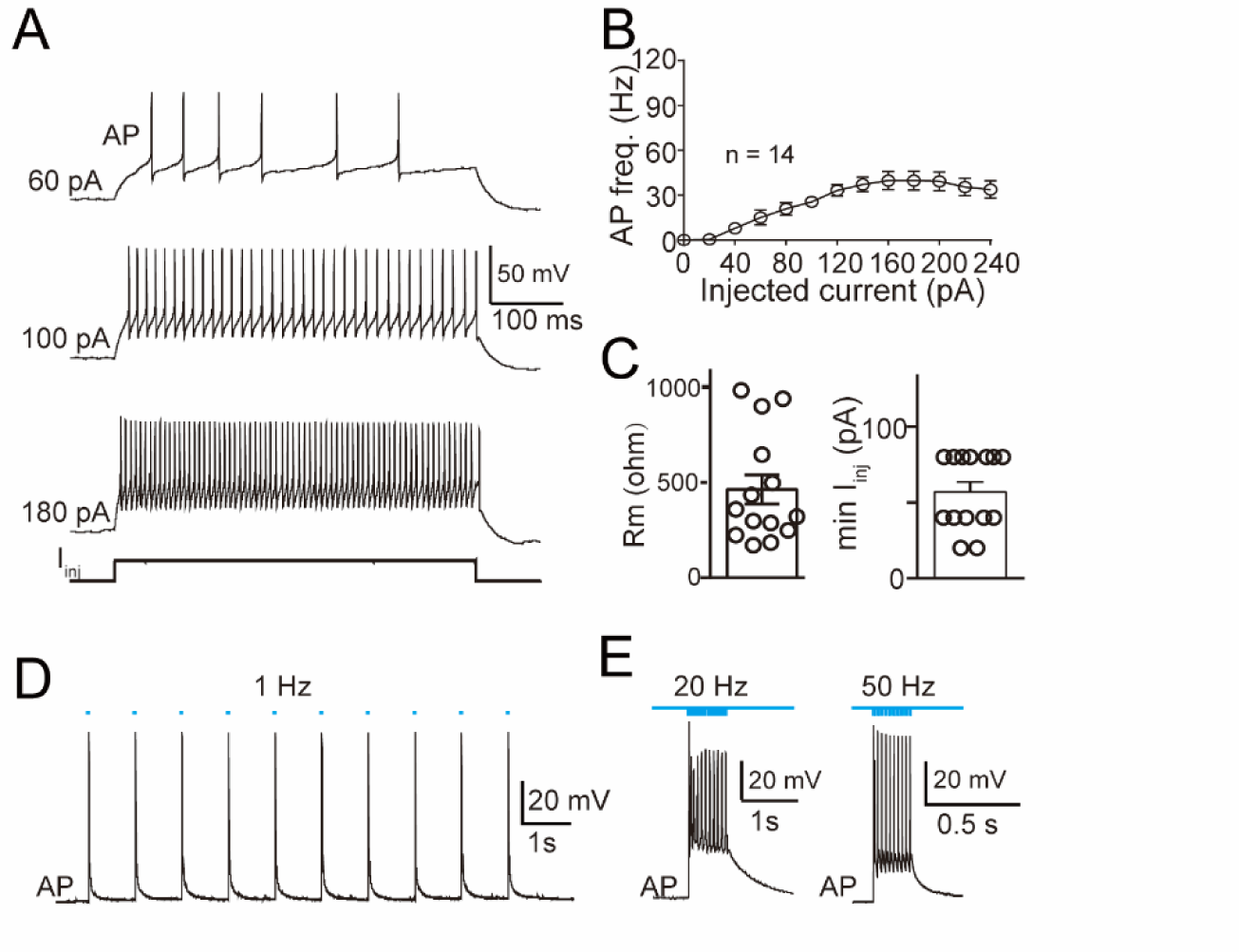
Functional properties of excitatory neurons in the SP5C. (A) Representative action potenial (AP) traces of Sp5 neurons evoked by current injection at increasing intensity, showing the neuron’s capability for firing APs at high frequencies. (B) Summary of AP firing frequency of SP5 neruons by injecting currents of increasing intensity. (C) Summary of cell membrane resistance (left) and minimum currents required for evoking AP for SP5 neurons (right). (D) Representative AP traces of Sp5 neurons stimulated by 1 Hz 470 nm light stimulation. (E) Representative AP traces of Sp5 neurons stimulated by 20 Hz (left) and 50 Hz (right) 470 nm light pulses, indicating the reliability of neuronal firing in response to high-frequency optogenenetic stimulation.

**Figure 5-Figure supplement 2.**
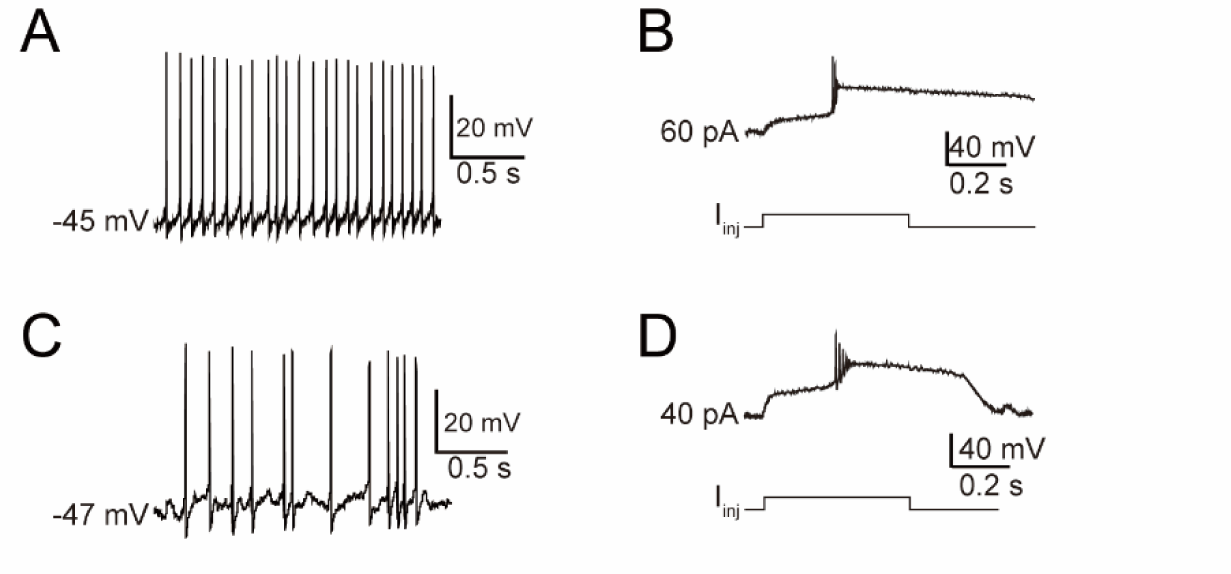
NaChB-overexpressing neurons exhibit spontaneous action potentials and sustained depolarization to injected currents. (A-B) Representative SP5 neuron overexpressing NaChBac fired spontaneous APs at high frequency (A) and evoked APs by injected current at the threshold level(B). (C-D) Another SP5 neuron overexpressing NaChBac fired spontaneous APs but at a revlatively low frequency (C) and evoked APs by injected current at the threshold level (D).

**Figure 5-Figure supplement 3.**
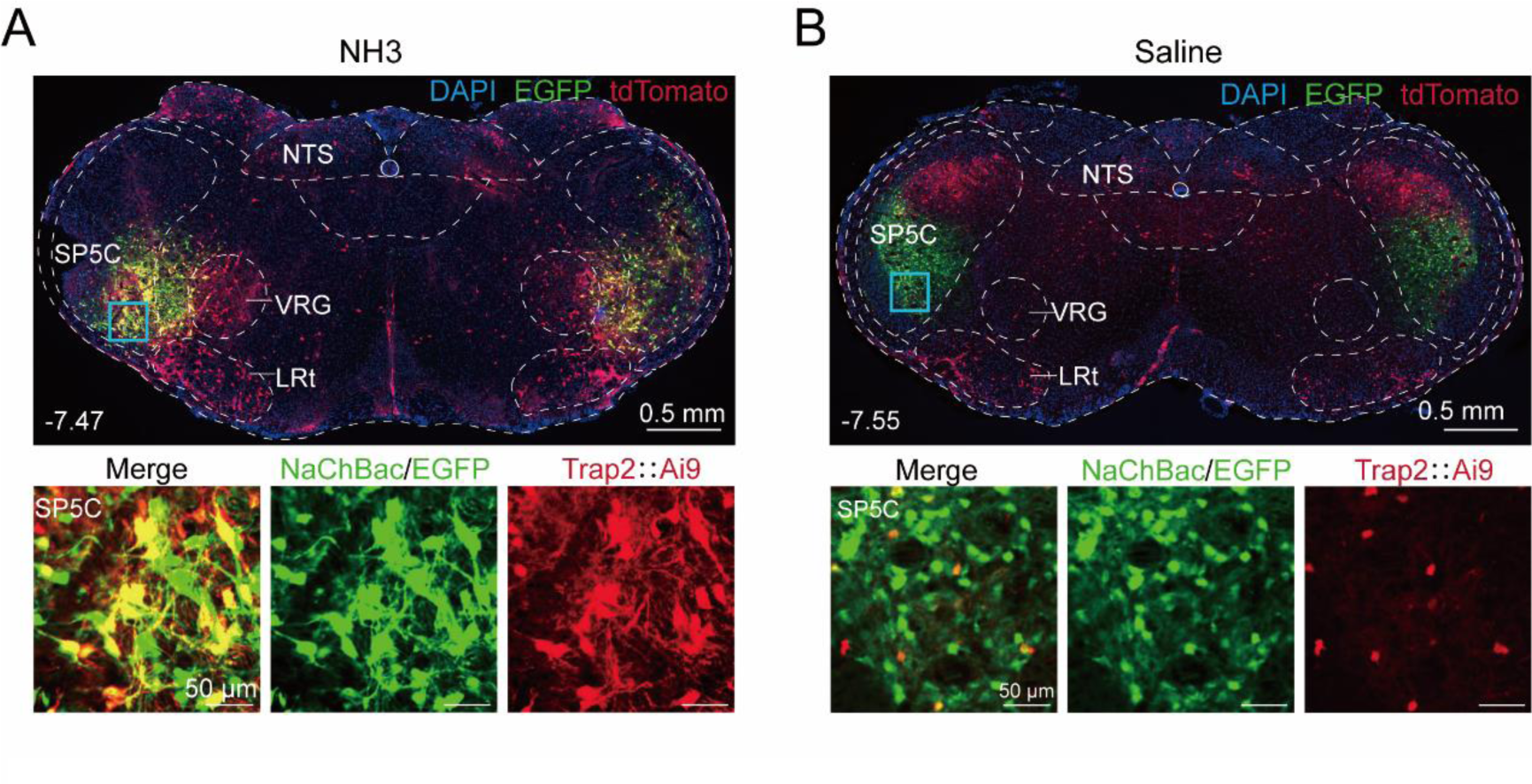
Overexpression of NaChBac enhances neuronal activity in the SP5C following NH3 nebulization. (A) Top, representative image showing TRAP2-labelled neurons after nebulized NH3 in TRAP2::Ai9 mice overexpressing NaChBac in the SP5C. Bottom, Enlarged region of interest highlighted by the blue rectangle in the Top image. (B) Top, representative image showing TRAP2-labelled neurons after nebulized saline in TRAP2::Ai9 mice overexpressing NaChBac in the SP5C. Bottom, Enlarged region of interest highlighted by the blue rectangle in the Top image.

## References

Allen, W.E., DeNardo, L.A., Chen, M.Z., Liu, C.D., Loh, K.M., Fenno, L.E., Ramakrishnan, C., Deisseroth, K., and Luo, L. (2017). Thirst-associated preoptic neurons encode an aversive motivational drive. Science 357, 1149–1155. 10.1126/science.aan6747.

Bianchi, A.L., and Gestreau, C. (2009). The brainstem respiratory network: an overview of a half century of research. Respir Physiol Neurobiol 168, 4–12. 10.1016/j.resp.2009.04.019.

Bolser, D.C. (2004). Experimental models and mechanisms of enhanced coughing. Pulm Pharmacol Ther 17, 383–388. 10.1016/j.pupt.2004.09.016.

Bolser, D.C., and Davenport, P.W. (2002). Functional organization of the central cough generation mechanism. Pulm Pharmacol Ther 15, 221–225. 10.1006/pupt.2002.0361.

Bongianni, F., Mutolo, D., Nardone, F., and Pantaleo, T. (2005). Ionotropic glutamate receptors mediate excitatory drive to caudal medullary expiratory neurons in the rabbit. Brain Res 1056, 145–157. 10.1016/j.brainres.2005.07.019.

Canning, B.J. (2006). Anatomy and neurophysiology of the cough reflex: ACCP evidence-based clinical practice guidelines. Chest 129, 33S–47S. 10.1378/chest.129.1_suppl.33S.

Canning, B.J. (2008). The cough reflex in animals: relevance to human cough research. Lung 186 Suppl 1, S23–28. 10.1007/s00408-007-9054-6.

Canning, B.J., Chang, A.B., Bolser, D.C., Smith, J.A., Mazzone, S.B., McGarvey, L., and Panel, C.E.C. (2014). Anatomy and neurophysiology of cough: CHEST Guideline and Expert Panel report. Chest 146, 1633–1648. 10.1378/chest.14-1481.

Canning, B.J., Mazzone, S.B., Meeker, S.N., Mori, N., Reynolds, S.M., and Undem, B.J. (2004). Identification of the tracheal and laryngeal afferent neurones mediating cough in anaesthetized guinea-pigs. J Physiol 557, 543–558. 10.1113/jphysiol.2003.057885.

Chang, R.B., Strochlic, D.E., Williams, E.K., Umans, B.D., and Liberles, S.D. (2015). Vagal Sensory Neuron Subtypes that Differentially Control Breathing. Cell 161, 622–633. 10.1016/j.cell.2015.03.022.

Chen, L.Y., Jiang, M., Zhang, B., Gokce, O., and Sudhof, T.C. (2017). Conditional Deletion of All Neurexins Defines Diversity of Essential Synaptic Organizer Functions for Neurexins. Neuron 94, 611–625 e614. 10.1016/j.neuron.2017.04.011.

Chen, Z., Lin, M.T., Zhan, C., Zhong, N.S., Mu, D., Lai, K.F., and Liu, M.J. (2022). A descending pathway emanating from the periaqueductal gray mediates the development of cough-like hypersensitivity. iScience 25, 103641. 10.1016/j.isci.2021.103641.

Chung, K.F., McGarvey, L., Song, W.J., Chang, A.B., Lai, K., Canning, B.J., Birring, S.S., Smith, J.A., and Mazzone, S.B. (2022). Cough hypersensitivity and chronic cough. Nat Rev Dis Primers 8, 45. 10.1038/s41572-022-00370-w.

Cinelli, E., Iovino, L., Bongianni, F., Pantaleo, T., and Mutolo, D. (2020). Essential Role of the cVRG in the Generation of Both the Expiratory and Inspiratory Components of the Cough Reflex. Physiol Res 69, S19–S27. 10.33549/physiolres.934396.

Coleridge, H.M., and Coleridge, J.C. (1994). Pulmonary reflexes: neural mechanisms of pulmonary defense. Annu Rev Physiol 56, 69–91. 10.1146/annurev.ph.56.030194.000441.

Cortese, A., Simone, R., Sullivan, R., Vandrovcova, J., Tariq, H., Yau, W.Y., Humphrey, J., Jaunmuktane, Z., Sivakumar, P., Polke, J., et al. (2019). Biallelic expansion of an intronic repeat in RFC1 is a common cause of late-onset ataxia. Nat Genet 51, 649–658. 10.1038/s41588-019-0372-4.

Del Negro, C.A., Funk, G.D., and Feldman, J.L. (2018). Breathing matters. Nat Rev Neurosci 19, 351–367. 10.1038/s41583-018-0003-6.

DeNardo, L.A., Liu, C.D., Allen, W.E., Adams, E.L., Friedmann, D., Fu, L., Guenthner, C.J., Tessier-Lavigne, M., and Luo, L. (2019). Temporal evolution of cortical ensembles promoting remote memory retrieval. Nat Neurosci 22, 460–469. 10.1038/s41593-018-0318-7.

Dhaka, A., Uzzell, V., Dubin, A.E., Mathur, J., Petrus, M., Bandell, M., and Patapoutian, A. (2009). TRPV1 is activated by both acidic and basic pH. J Neurosci 29, 153–158. 10.1523/JNEUROSCI.4901-08.2009.

Gannot, N., Li, X., Phillips, C.D., Ozel, A.B., Uchima Koecklin, K.H., Lloyd, J.P., Zhang, L., Emery, K., Stern, T., Li, J.Z., and Li, P. (2024). A vagal-brainstem interoceptive circuit for cough-like defensive behaviors in mice. Nat Neurosci. 10.1038/s41593-024-01712-5.

Guilleminault, L., Chazelas, P., Melloni, B., Magdelaine, C., Villeneuve, T., Brouquieres, D., Lia, A.S., and Magy, L. (2023). Repeat Expansions of RFC1 in Refractory Chronic Cough: A Missing Piece of the Puzzle? Chest 163, 911–915. 10.1016/j.chest.2022.11.014.

Kim, S.H., Hadley, S.H., Maddison, M., Patil, M., Cha, B., Kollarik, M., and Taylor-Clark, T.E. (2020). Mapping of Sensory Nerve Subsets within the Vagal Ganglia and the Brainstem Using Reporter Mice for Pirt, TRPV1, 5-HT3, and Tac1 Expression. eNeuro 7. 10.1523/ENEURO.0494-19.2020.

Klapoetke, N.C., Murata, Y., Kim, S.S., Pulver, S.R., Birdsey-Benson, A., Cho, Y.K., Morimoto, T.K., Chuong, A.S., Carpenter, E.J., Tian, Z., et al. (2014). Independent optical excitation of distinct neural populations. Nat Methods 11, 338–346. 10.1038/nmeth.2836.

Krohn, F., Novello, M., van der Giessen, R.S., De Zeeuw, C.I., Pel, J.J.M., and Bosman, L.W.J. (2023). The integrated brain network that controls respiration. Elife 12. 10.7554/eLife.83654.

Kupari, J., Haring, M., Agirre, E., Castelo-Branco, G., and Ernfors, P. (2019). An Atlas of Vagal Sensory Neurons and Their Molecular Specialization. Cell Rep 27, 2508–2523 e2504. 10.1016/j.celrep.2019.04.096.

Li, F., Jiang, H., Shen, X., Yang, W., Guo, C., Wang, Z., Xiao, M., Cui, L., Luo, W., Kim, B.S., et al. (2021). Sneezing reflex is mediated by a peptidergic pathway from nose to brainstem. Cell 184, 3762–3773 e3710. 10.1016/j.cell.2021.05.017.

Luo, F., Sclip, A., Jiang, M., and Sudhof, T.C. (2020). Neurexins cluster Ca(2+) channels within the presynaptic active zone. EMBO J 39, e103208. 10.15252/embj.2019103208.

Mazzone, S.B., and Undem, B.J. (2016). Vagal Afferent Innervation of the Airways in Health and Disease. Physiol Rev 96, 975–1024. 10.1152/physrev.00039.2015.

McGovern, A.E., Driessen, A.K., Simmons, D.G., Powell, J., Davis-Poynter, N., Farrell, M.J., and Mazzone, S.B. (2015). Distinct brainstem and forebrain circuits receiving tracheal sensory neuron inputs revealed using a novel conditional anterograde transsynaptic viral tracing system. J Neurosci 35, 7041–7055. 10.1523/JNEUROSCI.5128-14.2015.

Mutolo, D. (2017). Brainstem mechanisms underlying the cough reflex and its regulation. Respir Physiol Neurobiol 243, 60–76. 10.1016/j.resp.2017.05.008.

Nagai, Y., Miyakawa, N., Takuwa, H., Hori, Y., Oyama, K., Ji, B., Takahashi, M., Huang, X.P., Slocum, S.T., DiBerto, J.F., et al. (2020). Deschloroclozapine, a potent and selective chemogenetic actuator enables rapid neuronal and behavioral modulations in mice and monkeys. Nat Neurosci 23, 1157–1167. 10.1038/s41593-020-0661-3.

Park, J., Choi, S., Takatoh, J., Zhao, S., Harrahill, A., Han, B.X., and Wang, F. (2024). Brainstem control of vocalization and its coordination with respiration. Science 383, eadi8081. 10.1126/science.adi8081.

Prescott, S.L., Umans, B.D., Williams, E.K., Brust, R.D., and Liberles, S.D. (2020). An Airway Protection Program Revealed by Sweeping Genetic Control of Vagal Afferents. Cell 181, 574–589 e514. 10.1016/j.cell.2020.03.004.

Ren, D., Navarro, B., Xu, H., Yue, L., Shi, Q., and Clapham, D.E. (2001). A prokaryotic voltage-gated sodium channel. Science 294, 2372–2375. 10.1126/science.1065635.

Smith, J.C., Abdala, A.P., Borgmann, A., Rybak, I.A., and Paton, J.F. (2013). Brainstem respiratory networks: building blocks and microcircuits. Trends Neurosci 36, 152–162. 10.1016/j.tins.2012.11.004.

Su, Y., Barr, J., Jaquish, A., Xu, J., Verheyden, J.M., and Sun, X. (2022). Identification of lung innervating sensory neurons and their target specificity. Am J Physiol Lung Cell Mol Physiol 322, L50–L63. 10.1152/ajplung.00376.2021.

Subramanian, H.H., and Holstege, G. (2009). The nucleus retroambiguus control of respiration. J Neurosci 29, 3824–3832. 10.1523/JNEUROSCI.0607-09.2009.

Sun, H., Lin, A.H., Ru, F., Patil, M.J., Meeker, S., Lee, L.Y., and Undem, B.J. (2019). KCNQ/M-channels regulate mouse vagal bronchopulmonary C-fiber excitability and cough sensitivity. JCI Insight 4. 10.1172/jci.insight.124467.

Turner, R.D., Hirons, B., Cortese, A., and Birring, S.S. (2023). Chronic Cough as a Genetic Neurological Disorder? Insights from Cerebellar Ataxia with Neuropathy and Vestibular Areflexia Syndrome (CANVAS). Lung 201, 511–519. 10.1007/s00408-023-00660-4.

Vong, L., Ye, C., Yang, Z., Choi, B., Chua, S., Jr., and Lowell, B.B. (2011). Leptin action on GABAergic neurons prevents obesity and reduces inhibitory tone to POMC neurons. Neuron 71, 142–154. 10.1016/j.neuron.2011.05.028.

Wickersham, I.R., Lyon, D.C., Barnard, R.J., Mori, T., Finke, S., Conzelmann, K.K., Young, J.A., and Callaway, E.M. (2007). Monosynaptic restriction of transsynaptic tracing from single, genetically targeted neurons. Neuron 53, 639–647. 10.1016/j.neuron.2007.01.033.

Widdicombe, J.G. (1995). Neurophysiology of the cough reflex. Eur Respir J 8, 1193–1202. 10.1183/09031936.95.08071193.

Xie, Z., Li, D., Cheng, X., Pei, Q., Gu, H., Tao, T., Huang, M., Shang, C., Geng, D., Zhao, M., et al. (2022). A brain-to-spinal sensorimotor loop for repetitive self-grooming. Neuron 110, 874–890 e877. 10.1016/j.neuron.2021.11.028.

Xu, W., Morishita, W., Buckmaster, P.S., Pang, Z.P., Malenka, R.C., and Sudhof, T.C. (2012). Distinct neuronal coding schemes in memory revealed by selective erasure of fast synchronous synaptic transmission. Neuron 73, 990–1001. 10.1016/j.neuron.2011.12.036.

Xue, M., Atallah, B.V., and Scanziani, M. (2014). Equalizing excitation-inhibition ratios across visual cortical neurons. Nature 511, 596–600. 10.1038/nature13321.

Xue, Y., Mo, S., Li, Y., Cao, Y., Xu, X., and Xie, Q. (2024). Dissecting neural circuits from rostral ventromedial medulla to spinal trigeminal nucleus bidirectionally modulating craniofacial mechanical sensitivity. Prog Neurobiol 232, 102561. 10.1016/j.pneurobio.2023.102561.

Zhang, C., Lin, R.L., Hong, J., Khosravi, M., and Lee, L.Y. (2017). Cough and expiration reflexes elicited by inhaled irritant gases are intensified in ovalbumin-sensitized mice. Am J Physiol Regul Integr Comp Physiol 312, R718–R726. 10.1152/ajpregu.00444.2016.

